# Biparental vertical transmission of *Aedes* anphevirus, Guadeloupe mosquito virus, and verdadero virus in colonized *Aedes aegypti*

**DOI:** 10.64898/2026.07.20.739548

**Authors:** Tillie J. Dunham, Karla Saavedra-Rodriguez, Brian D. Foy, Christie E. Mayo, Mark D. Stenglein

## Abstract

*Aedes aegypti* mosquitoes can be infected by arboviruses that can be transmitted to and cause disease in vertebrates. *Ae. aegypti* can also be infected by insect-specific viruses, which do not infect vertebrates, but may impact mosquito biology and vector competence. In this study we characterized the viromes of *Ae. aegypti* populations that had been maintained in laboratory colonies for 4 to 18 years. Mosquitoes were originally collected in Tapachula, Poza Rica and Merida, Mexico and New Orleans, USA. We used metagenomics to characterize the viruses present in the colonies, then quantified the vertical transmission efficiencies of the viruses by performing crosses between populations. The viruses infecting these colonies included *Aedes* anphevirus, Guadeloupe mosquito virus, and verdadero virus. These viruses were at high prevalence in infected populations, with over 75% of individual adult mosquitoes infected. All three viruses exhibited biparental vertical transmission: both infected mothers and infected fathers transmitted infection to their offspring, but in all cases maternal transmission was more efficient than paternal. We also identified evidence of possible horizontal transmission between adult mosquitoes that had cohabited during crosses. Efficient transgenerational transmission likely contributes to the ability of these viruses to persist in laboratory populations and nature. Because of their efficient vertical transmission and minimal apparent fitness costs, these viruses could be good candidates for gene delivery to mosquito populations, including for vector control.

**Author summary:** *Aedes aegypti* mosquitoes are responsible for transmission of many dangerous pathogens, including dengue virus, Zika virus, and yellow fever virus. These mosquitoes also carry viruses that do not pose a threat to public health, but might influence mosquito biology. In this study, we identified viruses that persistently infected 4 long-established *Ae. aegypti* colonies in our insectary. These viruses were at high prevalence in these laboratory populations, and did not appear to harm infected mosquitoes. The viruses exhibited an ability to efficiently transmit from parents to offspring, both from infected mothers and infected fathers. This provided an explanation for how these mosquitoes were able to persist so successfully in laboratory populations, and by extension in nature. Given their ability to efficiently spread and minimal apparent fitness costs, these viruses represent promising candidates for use in mosquito control or other practical applications.

## Introduction

*Aedes aegypti* mosquitoes transmit important arbovirus pathogens including dengue, Zika, and chikungunya viruses (1). Successful transmission of arboviruses depends on their ability to infect, disseminate within, and ultimately reach the salivary glands of the mosquito, where they can be transmitted to vertebrate hosts during blood feeding (2). Although the global burden of arboviral disease is high, only a small fraction of mosquitoes in natural populations are infected by arboviruses at any given time (3). In contrast, mosquitoes are commonly infected by so-called insect-specific viruses (ISVs) (4–9). ISVs have no apparent ability to infect vertebrates and few well characterized fitness impacts on their mosquito hosts. A growing number of studies have characterized the virome of wild mosquitoes (10–20). Mosquito-infecting ISVs are diverse in terms of their prevalence and genetics, with representatives from all major groups of viruses, and are increasingly recognized as important components of mosquito biology.

ISVs are relevant to vector-borne diseases because they could naturally alter vector competence or could be used to deliver genes to mosquito populations (5, 6, 21–23). There is experimental and observational evidence that ISVs can interfere with arbovirus replication in cultured cells or reduce vector competence (24–32). Nevertheless, arboviruses remain a substantial public health problem despite the ubiquity of ISVs, and the impact of ISVs on vector competence is likely a function of complex environmental and genetic variables. In addition, ISVs could be genetically engineered to block arbovirus replication beyond their natural ability to do so (21, 22). Alternatively, ISVs, especially common ones, could provide a means to study mosquito population structure and ecology (33).

Established laboratory colonies provide a useful system for studying ISV persistence. The virome of lab-colonized mosquitoes appears to be less diverse than that of their wild counterparts, with certain ISVs tending to predominate (16, 34–36). (35, 36). These viruses presumably have properties that enable them to persist in laboratory populations, including efficient transmission and minimal fitness costs.

Here, we used metagenomic sequencing to characterize the virome of four colonies of *Ae. aegypti* with distinct geographic origins maintained in our insectary. We performed experimental crosses to investigate transmission from infected mothers or fathers to offspring and evaluated evidence of possible horizontal transmission. Inclusion of distinct outbred populations in experiments enabled us to assess whether transmission efficiency might vary as a function of mosquito genotype. This offered insight into mechanisms that promote long-term viral persistence in mosquito populations.

## Materials and Methods

### Experimental overview

The study consisted of two phases. First, we used metagenomic sequencing to identify ISVs persisting in four laboratory colonies of *Ae. aegypti* that originated from different geographic locations. Second, we quantified transmission efficiencies using controlled crosses and RT-qPCR screening of individual mosquitoes. Crosses between infected and uninfected colonies were designed to distinguish maternal and paternal transmission, while screening of previously uninfected adults after cohabitation provided evidence for potential horizontal transmission. Transmission efficiencies were estimated as the proportion of offspring or exposed adults testing positive for each virus.

### Mosquito colonies

All colonies were maintained in the insectary of the Center for Vector-Borne Infectious Diseases at Colorado State University in Fort Collins, Colorado, USA. Poza Rica *Ae. aegypti* were originally collected from Poza Rica, Veracruz, Mexico in 2012 (37) (**Supplemental Figure 1**). Tapachula *Ae. aegypti* were originally collected from Tapachula, Chiapas, Mexico in 2018 (38). New Orleans *Ae. aegypti* were originally collected in New Orleans, Louisiana, USA in 2005 (39). Vergel *Ae. aegypti* were originally collected in Mérida, Yucatán, Mexico in 2011 (40). Experiments were performed in 2022.

### Mosquito rearing

At all life stages, mosquitoes were housed at 27°C and 75-80% humidity. Eggs were hatched in 5×12 inch plastic bins with 800 mL of autoclaved room temperature deionized water (DI H2O). Growth containers were covered with organdy fabric secured by rubber bands. Egg papers were removed 24 hours after hatching and larvae were moved into 16×32 inch plastic containers with 2 inches of tap water. Aquatic stages were provided with 1/4 teaspoon of ground TetraMin fish food (Tetra). Larval crowding was checked daily, splitting the bin as necessary until pupation began, ∼7 days after hatching. Pupae were picked with a sterile Pasteur pipette or mesh scoop and placed into a small cup with water to be sorted by sex. Female and male pupae were housed separately in 64 oz paper cartons until they reached adulthood. Adults were provided with raisins daily as a source of sugar.

### Experimental crosses

Crosses were made by combining 80-100 virgin females with 15-30 virgin males, derived from pupae that had been separated by sex. Mosquitoes were anesthetized at 4 °C for 5 minutes, counted and then placed into a new 64 oz paper carton with an organdy screen on top. Adults were provided with sterile water and raisins until blood feeding. Crosses were performed in duplicate and involved combinations of male and female adults from the different colonies.

### Artificial blood feeding

Prior to blood feeding, mosquitoes were deprived of raisins for 24 hours and water for 3-6 hours. 4-day old female mosquitoes were provided 2 mL of warm defibrinated calf blood (Colorado Serum Company) via an artificial membrane feeding system. The artificial membrane feeding system (Lillie Glassblowers) has a glass blood feeder covered with hog’s gut and was kept at 37°C using a circulating water bath. Females were allowed to feed for 1 hour. After feeding, females were transferred to cartons for egg collection.

### Egg collection

Egg collection containers were made with a small water cup lined with a thin strip of paper towel (egg paper) sitting in sterile DI H2O. Egg collection containers were taped to the bottom of cartons containing blood fed adults. 5 days after feeding, egg papers were allowed to dry completely at 27°C and 75-80% humidity.

### Total RNA Extraction

Individual adult mosquitoes were placed in wells of a 96-well deep-well plate (Costar, 3958) with 1 ball bearing (McMaster-Carr 1598K22) and 100 µL of lysis buffer containing 5M guanidine thiocyanate, 0.1M Tris(hydroxymethyl)aminomethane (Tris), pH7.5, 0.01M ethylenediaminetetraacetic acid (EDTA), and 20 mM dithiothreitol (DTT). Mosquitoes were homogenized in a Qiagen TissueLyzer instrument at 30 Hz for 3 minutes. After homogenization, samples were transferred to a 96-well deep well KingFisher Plate and mixed with 90 µL of washed Sera-Mag Speedbead magnetic beads (Cytiva, 65152105050250), 60 µL of 100% isopropanol, and 10 µL of lysis binding enhancer containing 200 µg/mL of proteinase K (P8107S, New England Biolabs), 20% glycerol, and 0.5% sodium dodecyl sulfate (SDS). RNA was purified using a KingFisher instrument with two wash steps. The first wash buffer contained 10 mM Tris, pH 7.5, 900 mM guanidine thiocyanate, and 20% ethanol. The second wash buffer consisted of 10 mM Tris, pH8, 1 mM EDTA, and 80% ethanol. RNA was eluted into 60 µL of water. Purified RNA was stored at −80°C until further analysis.

### Metagenomic sequencing

Total RNA from 12 male and 12 female adults from each colony were pooled and used as input for library preparation. Libraries were constructed using the Kapa Biosystems RNA Hyper Prep kit following the manufacturer’s protocol (Roche). Library size distribution was assessed using an Agilent tapestation instrument. Libraries were sequenced on a NovaSeq X Plus instrument using paired-end 2×150 sequencing (Azenta Life Sciences). We included a negative control library prepared from water and a positive control library prepared from HeLa cell total RNA.

Virus sequences were recovered from metagenomic datasets as previously described (41). Briefly, low quality and adapter-derived bases were trimmed from reads using Cutadapt v3.5 (42). Host-derived reads were removed by mapping trimmed reads to the *Ae. aegypti* genome and transcriptome AaegL5.0 assembly (GCF_002204515.2) using bowtie2 v2.4.5 (43, 44). Remaining reads were assembled using the SPAdes assembler v3.15.4 (45). Virus-derived contigs were identified using BLASTN and BLASTX to search the National Center for Biotechnology Information (NCBI) nucleotide and protein databases (46). Virus sequences were validated by remapping unassembled reads to draft sequences using bowtie2 and annotated using Geneious software v2025.0.3 (https://www.geneious.com). Code is available at: https://github.com/stenglein-lab/taxonomy_pipeline and https://github.com/stenglein-lab/remapping_workflow.

### Detection and quantification of viral RNA

Virus-specific RT-qPCR assays were used to determine infection status and quantify relative viral RNA levels in individual mosquitoes. Assays targeted *Aedes* anphevirus, verdadero virus, and Guadeloupe mosquito virus Supplemental Table 1). cDNA was synthesized by adding 5.5 µL of RNA, 1 µL of a 250 µM random 15mer oligonucleotide, 1 µL of 10 mM each deoxynucleotide triphosphates (dNTPs; NEB) and 5.5 µL of water. Reactions were incubated at 65°C for 5 minutes then on ice for 1 minute. A mix containing 4 µL 5x first strand buffer (Invitrogen), 1 µL 0.1 M DTT, and 1 µL Superscript III reverse transcriptase (Invitrogen) was then added to each reaction. Reactions were incubated at 50°C for 60 minutes then 80°C for 10 minutes. Resulting cDNA was diluted with water to 100 µL total volume.

We used 2.5 µL of diluted cDNA as input to qPCR reactions containing 1x Luna qPCR Master Mix (NEB), 0.5 µM forward primer, 0.5 µM reverse primer (primer sequences in Supplemental Table 1), and water to a final volume of 10 µL. We performed qPCR on a QuantStudio3 instrument (ThermoFisher) using the following thermocycling conditions: 95°C for 3 minutes, 40 cycles of 95°C for 10 seconds, 60°C for 45 seconds, followed by a melting curve analysis. 12 females and 10-12 male mosquitoes were tested from each experimental group. 2 males from certain groups were replaced with a negative control (water) or a positive control (known infected mosquitoes from the Poza Rica colony). Legitimate target amplification was confirmed by inspection of melting curve peaks. Mosquitoes were classified as positive when amplification produced the expected melting temperature profile and we did not use a maximum Ct cutoff beyond which mosquitoes would be called negative. Primers targeting the host actin-5C mRNA were used as an internal positive control Supplemental Table 1 (47). Levels of viral RNA were normalized to levels of actin-5C mRNA. Testing involved a total of 4472 qPCR reactions.

### Endogenous Viral Element (EVE) testing

Because low-level GMV RT-qPCR signals were detected in colonies lacking detectable GMV reads in metagenomic datasets, we evaluated whether amplification could have resulted from transcription of a GMV endogenous viral element (EVE) integrated into the host genome. We performed qPCR as previously described, but without prior reverse transcription, on purified nucleic acid, which contained both RNA and DNA. Detection under these conditions would have indicated amplification from genomic DNA rather than viral RNA.

### Modeling of transmission efficiency

We performed logistic regression modeling to evaluate the impact of experimental variables on transmission efficiency. Maternal transmission efficiency was defined as the proportion of offspring testing positive following crosses between infected females and uninfected males. Paternal transmission efficiency was defined as the proportion of offspring testing positive following crosses between infected males and uninfected females. Potential horizontal transmission was assessed using the proportion of previously uninfected adults that tested positive following cohabitation and mating with infected mosquitoes. Because infection status of individual parents was not determined, estimates reflected population-level transmission efficiencies.

Models used the lme4 R package version 2.0-1 (48). Models for vertical transmission took the form: infected ∼ offspring_sex + non_infected_parent_colony (New Orleans vs Vergel) + infected_parent_sex (mother vs. father). Models for horizontal transmission took the form: infected ∼ exposed_parent_sex + exposed_parent_colony (New Orleans vs. Vergel). Models for Guadeloupe mosquito virus transmission also included a infected_parent_colony (Poza Rica vs. Tapachula) term. Infected was a binary variable indicating whether individual mosquitoes were qRT-PCR positive for the indicated virus. Fixed effects included the sex of tested offspring or cohabiting adult parent, the infected and non-infected parental colonies, and, for vertical transmission, the infected parent sex: infected mothers (maternal transmission), of infected fathers (paternal transmission). Models that included replicate as a random effect were not significantly better fitting as determined by ANOVA, so models shown include fixed effects only. An exception was horizontal transmission of Guadeloupe mosquito virus: a model that included replicate as a random effect fit significantly better. Models that included interaction terms were not significantly better fitting so models do not include interaction terms. Code for modeling is provided in the linked github repository.

### Data Availability

Data and code for all analyses are available at https://github.com/tdunham19/CM3_Mosquito_Paper. Assembled virus sequences are available in the NCBI nucleotide database under accessions PV646265-PV646272. Metagenomic datasets are available from the NCBI SRA database under bioproject PRJNA1260027. This paper is implemented as a reproducible quarto markdown document (49).

**Supplemental Figure 1:**
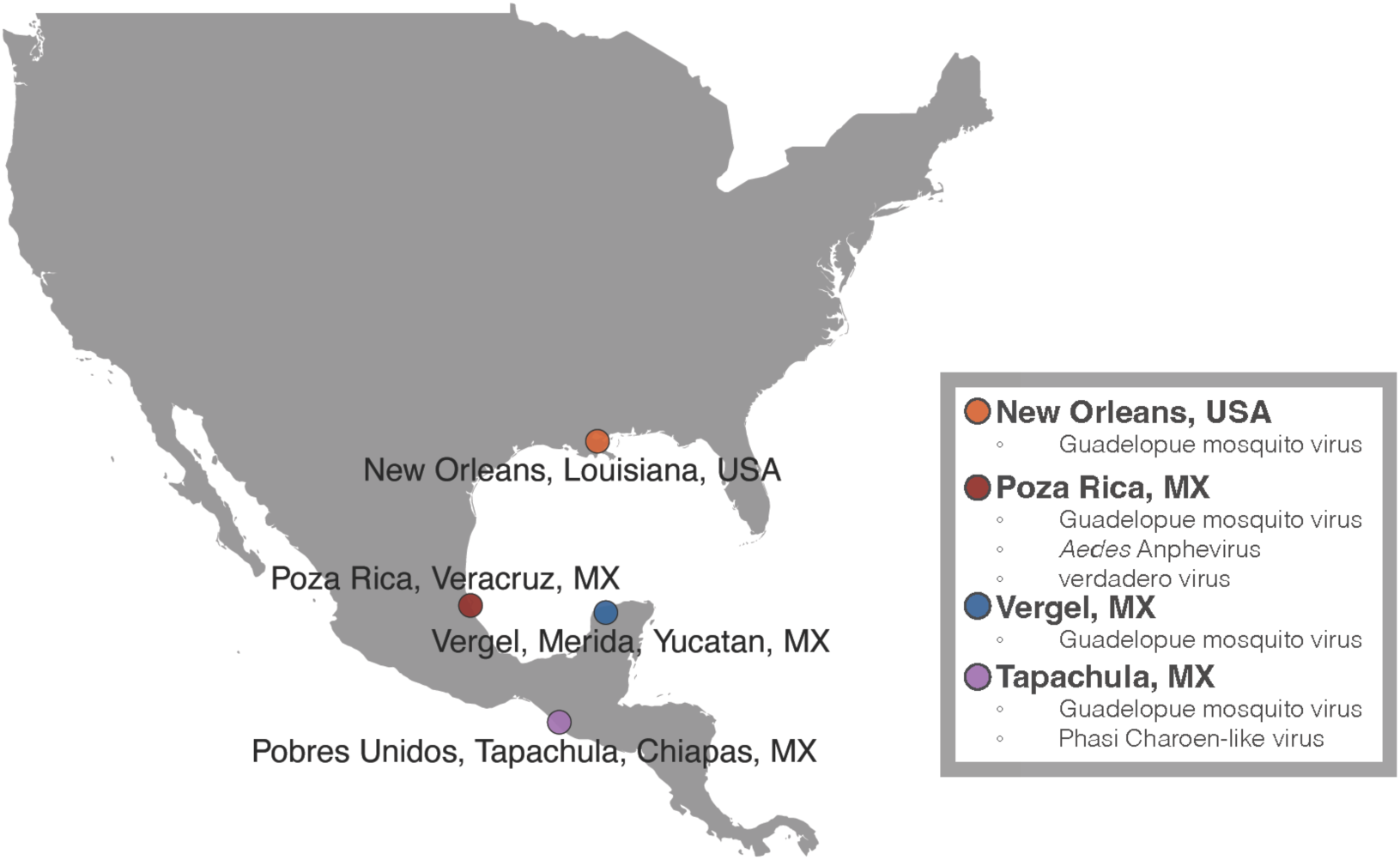
Map of original collection locations of Aedes aegypti colonies used for this project.

## Results

### Virome characterization of laboratory *Ae. aegypti* colonies

We used shotgun metagenomic sequencing to characterize the viruses infecting *Ae. aegypti* in four colonies maintained in our insectary. We prepared libraries from total RNA from mixed-sex pools of parental mosquitoes from experimental crosses. Libraries were sequenced on an Illumina NovaSeq X instrument to generate an average of 6.3×10^6^ read pairs per dataset. Following quality and adapter trimming and removal of host-mapping reads, an average of 4.1×10^5^ read pairs per dataset remained (6%). Remaining reads were assembled and contigs were used as BLAST queries to the NCBI nucleotide and protein databases to identify virus contigs.

This analysis yielded 8 contigs longer than 1000 nt corresponding to sequences from 4 viruses all known to infect *Ae. aegypti*: Guadeloupe mosquito virus (GMV) (17), verdadero virus (41), chaq-like virus, which is likely a satellite of verdadero virus (41), and *Aedes* Anphevirus (AeAV) (34) (**Table 1**). The Poza Rica colony produced high coverage coding complete sequences for AeAV, GMV, veradero virus, and chaq-like virus. These sequences were all >99% identical to existing sequences in the NCBI nucleotide database (**Table 1**; **Supplemental Figure 2**). The Tapachula dataset yielded a coding complete GMV sequence. Neither the New Orleans nor the Vergel colonies produced any candidate viral contigs longer than 1000 nt. Datasets contained shorter contigs with similarity to virus sequences that likely originated from the endogenized viral sequences that are common in *Aedes* genomes (50, 51). It is also possible that short contigs derived from low prevalence or low level viral infections, which we did not investigate further. No reads in the New Orleans or Vergel datasets mapped to the virus sequences detected in the Poza Rica or Tapachula colonies.

**Table 1:** Virus sequences identified in Aedes aegypti colonies using shotgun metagenomic sequencing. (a) The most similar existing sequence in the NCBI nucleotide database, determined using a BLASTN search. (b) Accessions of newly deposited sequences from this study. (c) The average depth of coverage in metagenomic datasets.

| Colony | Virus | Most similar Genbank sequence (a) | Identity to most similar sequence (%) | Genbank accession (b) | Average coverage depth (x) (c) |
| --- | --- | --- | --- | --- | --- |
| Poza Rica | Aedes anphevirus | MW435013 | 99.6 | PV646265 | 38x |
| Poza Rica | verdadero virus RNA 1 & 2 | MT742174, MT742175 | 99.8-99.9 | PV646269-PV646270 | 356x |
| Poza Rica | chaq-like virus | MT742176 | 100 | PV646266 | 910x |
| Poza Rica | Guadeloupe mosquito virus RNA 1 & 2 | MW434816, MW434807 | 99.3-99.6 | PV646267-PV646268 | 1202x |
| Tapachula | Guadeloupe mosquito virus RNA 1 & 2 | MW434816, MN053806 | 99.1-99.6 | PV646271-PV646272 | 589x |

In summary, metagenomic sequencing identified the sequences of four ISVs: *Aedes* anphevirus, verdadero virus, chaq-like virus, and Guadeloupe mosquito virus (GMV). These viruses were detected in the Poza Rica and Tapachula colonies but not the New Orleans and Vergel colonies. High depth of coverage suggested that these viruses occurred at high abundance and prevalence within infected colonies, so were the focus of subsequent transmission experiments.

**Supplemental Figure 2:**
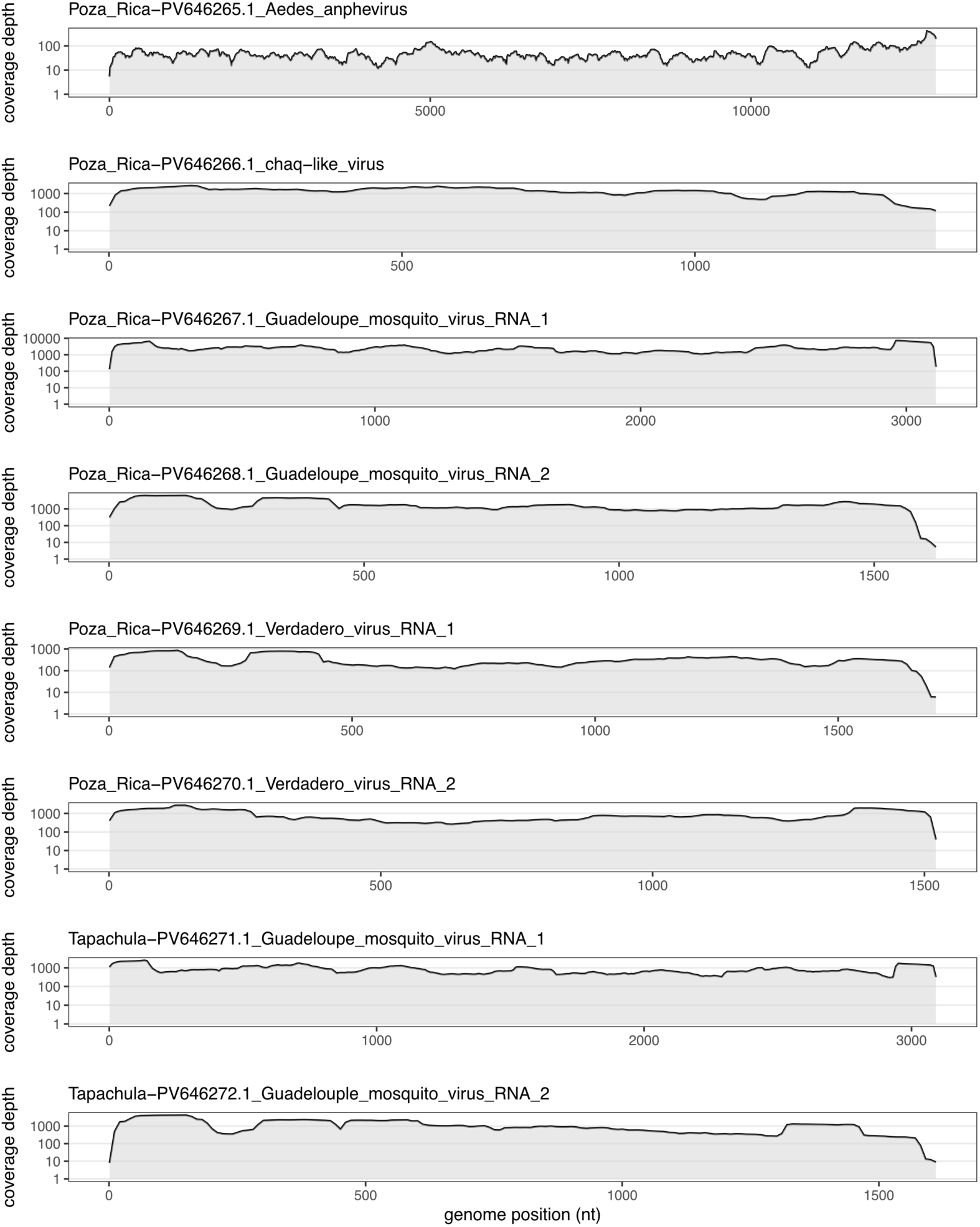
Coverage depth of virus sequences identified in Aedes aegypti colonies using shotgun metagenomic sequencing. Average coverage depth of mapped reads in 10-nucleotide windows is plotted. NCBI nucleotide accessions of virus sequences are indicated.

To determine the efficiency with which these viruses were transmitted from adults to offspring, we set up a series of crosses involving mosquitoes from the different colonies (**Supplemental Figure 1, Supplemental Figure 3**). We used RT-qPCR to quantify the fraction of infected individuals (prevalence) and viral RNA levels in individual mosquitoes. Sampling of adults at the time of crossing allowed us to measure prevalences and RNA levels in parental populations. Sampling of adults after cohabitation and mating provided an estimate of possible horizontal transmission. Sampling of adult offspring from crosses revealed the efficiency with which the viruses transmitted vertically from parents to the next generation (**Supplemental Figure 3**). By setting up crosses with males or females from the different populations we were able to separately quantify paternal and maternal transmission efficiencies and evaluate potential impacts of different parental genotypes. We performed each cross twice and report the results of both replicate experiments.

**Supplemental Figure 3:**
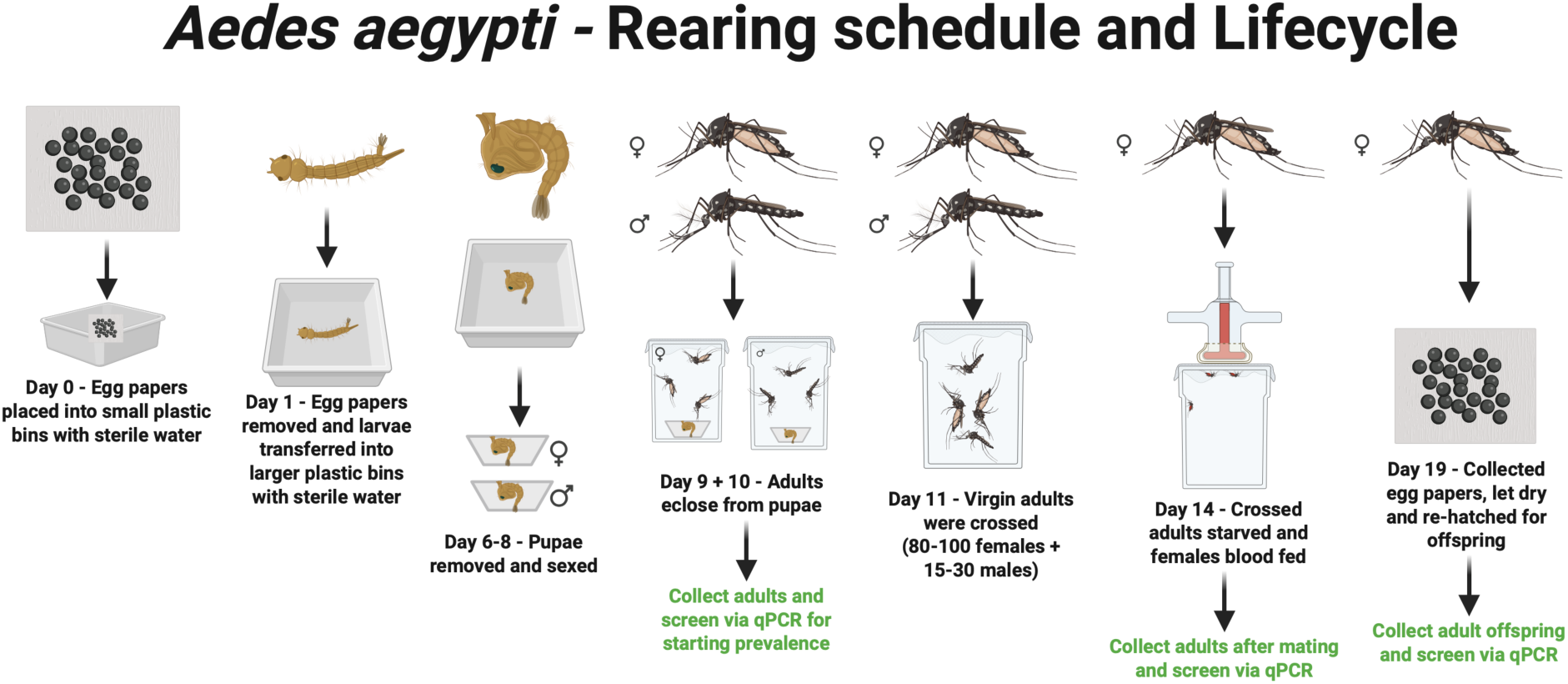
Aedes Aegypti Rearing Schedule and Lifecycle: Outline of experiment, Aedes aegypti life cycle and screening time points throughout the experiment. Created in BioRender. Dunham, T. (2026) https://BioRender.com/1wb7iq3.

### Aedes anphevirus

*Aedes* anphevirus is a member of the order *Mononegavirales*, which includes diverse monopartite negative-sense single-stranded RNA viruses (34, 52). Related viruses have been identified in other *Aedes* and *Anopheles* species (e.g., (12, 53)). The detection of anphevirus in *Ae. aegypti* sperm and eggs suggested that this virus could be vertically transmitted, potentially from either parent (34, 53).

Nearly all tested adult mosquitoes in the Poza Rica colony were anphevirus-infected: 12/12 (100%) of males and 11/12 (92%) of females were positive by RT-qPCR (**Figure 1**). The mean level of anphevirus RNA in infected males and females was 8.8× and 1.1× the level of actin mRNA, respectively. No mosquitoes in the New Orleans or Vergel colonies tested positive for AeAV by metagenomic sequencing or by RT-qPCR.

**Figure 1:**
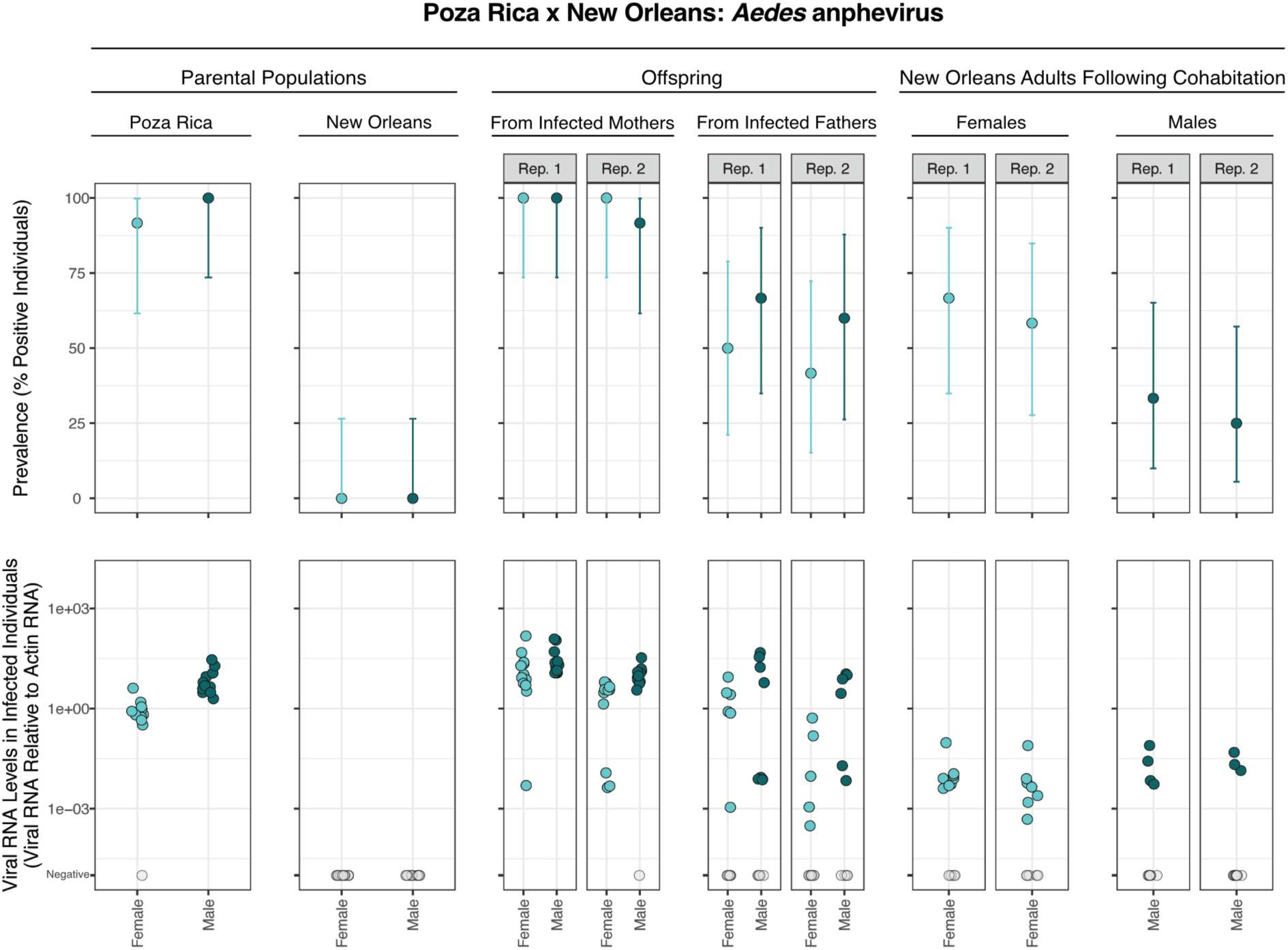
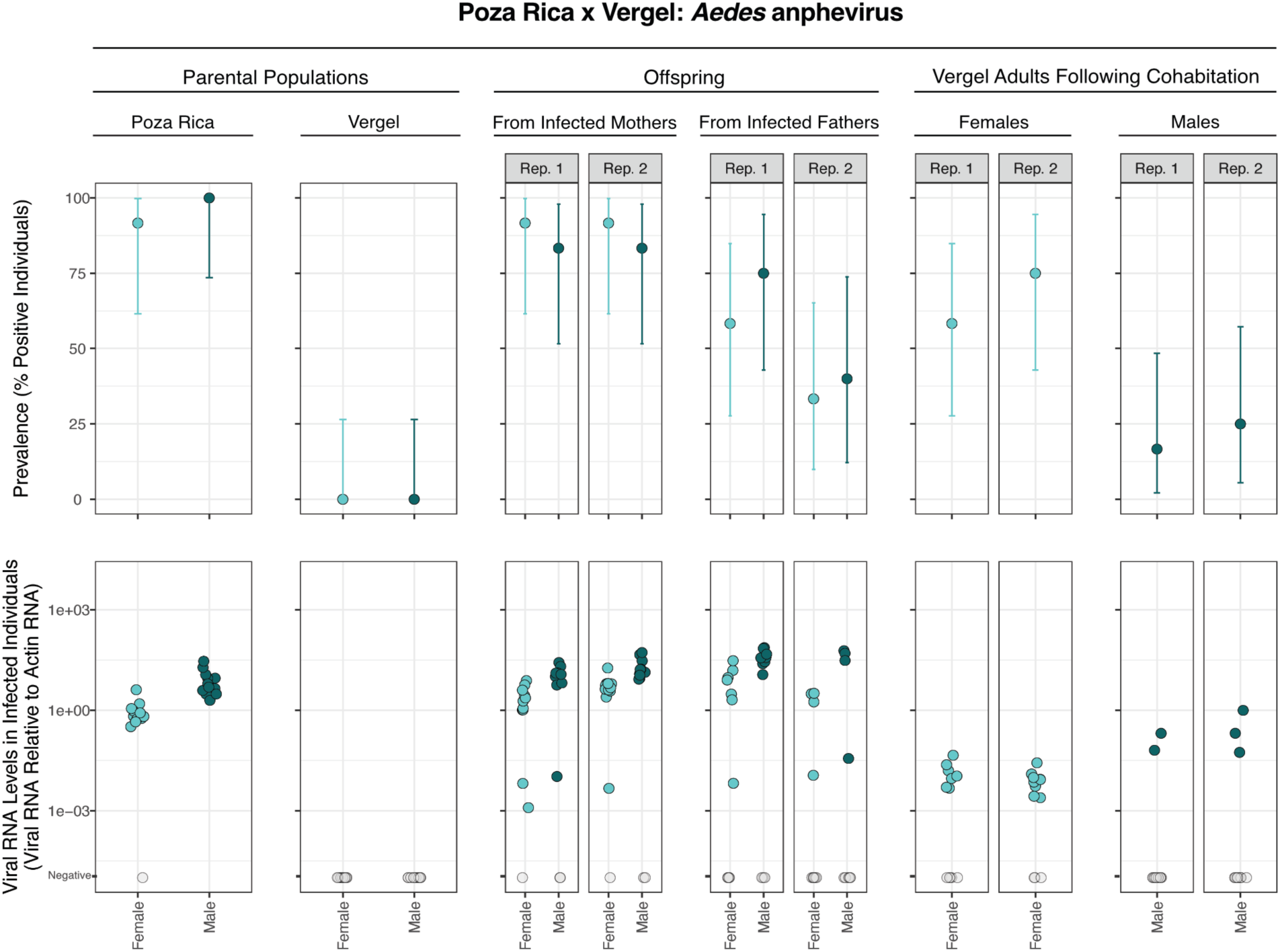
Prevalence and transmission of Aedes anphevirus. Upper panels show prevalence of AeAV in different populations of parental and offspring mosquitoes. Error bars indicate binomial 95% confidence intervals. Lower panels show levels of AeAV RNA detected by RT-qPCR in individual mosquitoes normalized to levels of actin mRNA. Values for uninfected mosquitoes are plotted in grey on the lowest Y axis position. Rep refers to replicate crosses. (A) Crosses between mosquitoes from the Poza Rica colony and the New Orleans colony. (B) Crosses between mosquitoes from the Poza Rica colony and the Vergel colony. The same data for parental Poza Rica is shown in upper and lower panels.

AeAV transmitted efficiently from infected mothers to offspring (**Figure 1**). 89/96 (93%) of the offspring of Poza Rica mothers tested positive for anphevirus RNA. Transmission from infected fathers was less efficient: 49/92 (53%) of the offspring of infected fathers tested positive. The mean level of anphevirus RNA in infected offspring was similar for maternal and paternal transmission: 15.0× and 15.1× the level of actin mRNA, respectively. Offspring exhibited a bimodal distribution of anphevirus RNA levels (**Figure 1**). A subset of positive offspring had relatively high anphevirus RNA levels (higher than actin mRNA levels), and another subset had levels ∼1000× lower. Transmission efficiencies were similar in crosses involving uninfected parents from the New Orleans and the Vergel colonies (**Figure 1**).

We used logistic regression to model anphevirus transmission (**Table 2**). Models for vertical transmission contained transmission mode (maternal vs. paternal), uninfected parent colony (New Orleans vs. Vergel), and offspring sex as fixed effects. Models that included replicate as a random effect were not significantly better fitting. For vertical transmission, transmission mode was the only significant predictor variable, with offspring more likely to be infected via maternal transmission (p = 5.5×10^−8^; **Table 2**).

**Table 2:** Logistic regression model of anphevirus transmission.

| Predictor | Vertical transmission |  |  | Horizontal transmission |  |  |
| --- | --- | --- | --- | --- | --- | --- |
|  | OR | 95% CI | p-value | OR | 95% CI | p-value |
| (Intercept) | 13.9 | 5.84, 38.7 | <0.001 | 1.92 | 0.93, 4.14 | 0.085 |
| Offspring or exposed parent sex [male] | 1.33 | 0.64, 2.78 | 0.4 | 0.18 | 0.07, 0.43 | **<0.001** |
| Non-infected parent colony [Vergel] | 0.66 | 0.32, 1.37 | 0.3 | 0.90 | 0.37, 2.18 | 0.8 |
| Vertical transmission mode [paternal] | 0.09 | 0.03, 0.20 | **<0.001** |  |  |  |
| No. Obs. | 188 |  |  | 96 |  |  |
| AIC | 183.5 |  |  | 122.3 |  |  |

Abbreviations: CI = Confidence Interval, OR = Odds Ratio

We tested previously uninfected parents after cohabitation and mating with infected parents to assess possible horizontal transmission. 12/48 (25%) of males and 31/48 (65%) of females were positive by RT-qPCR following cohabitation with opposite sex infected mosquitoes (**Figure 1**). However, levels of AeAV RNA in these mosquitoes were low, on average 0.09× levels of actin mRNA (166× lower than average levels in infected offspring). These positive signals may represent legitimate low-level infection following horizontal transmission or could have resulted from cross-contamination of viral RNA during cohabitation or mating.

Models for horizontal transmission included exposed parent colony (New Orleans vs Vergel) and sex as fixed effects. For horizontal transmission, exposed parent sex was the only significant predictor, with females more likely to test positive following exposure to infected fathers (p = 1.6×10^−4^; **Table 2**). The higher prevalence in females may reflect possible sexual transmission during mating.

### Verdadero virus

Verdadero virus is a two-segmented double-stranded (ds) RNA virus similar to viruses in the family *Partitiviridae* (41, 54). Poza Rica mosquitoes were infected by verdadero virus and by chaq-like virus, a presumed satellite virus of verdadero virus (41). We did not assess transmission of chaq-like virus separately. Previous experimental evidence showed that verdadero virus could be vertically transmitted with high efficiency (41). A related virus, Palmetto partiti-like virus, was found to be at high prevalence in colonized *Ae. aegypti* from Florida and capable of vertical transmission (36).

Verdadero virus was at high prevalence in the Poza Rica parental population, and undetectable in the New Orleans or Vergel mosquitoes (**Figure 2**). 11/12 (92%) of Poza Rica males and 10/12 (83%) of females were positive by RT-qPCR (**Figure 2**). The mean level of verdadero virus RNA 1 in infected males and females was 2.6× and 0.3× the level of actin mRNA, respectively. Galbut virus, a related partitivirus that infects *Drosophila melanogaster*, also exhibits higher average RNA levels in infected males (41).

**Figure 2:**
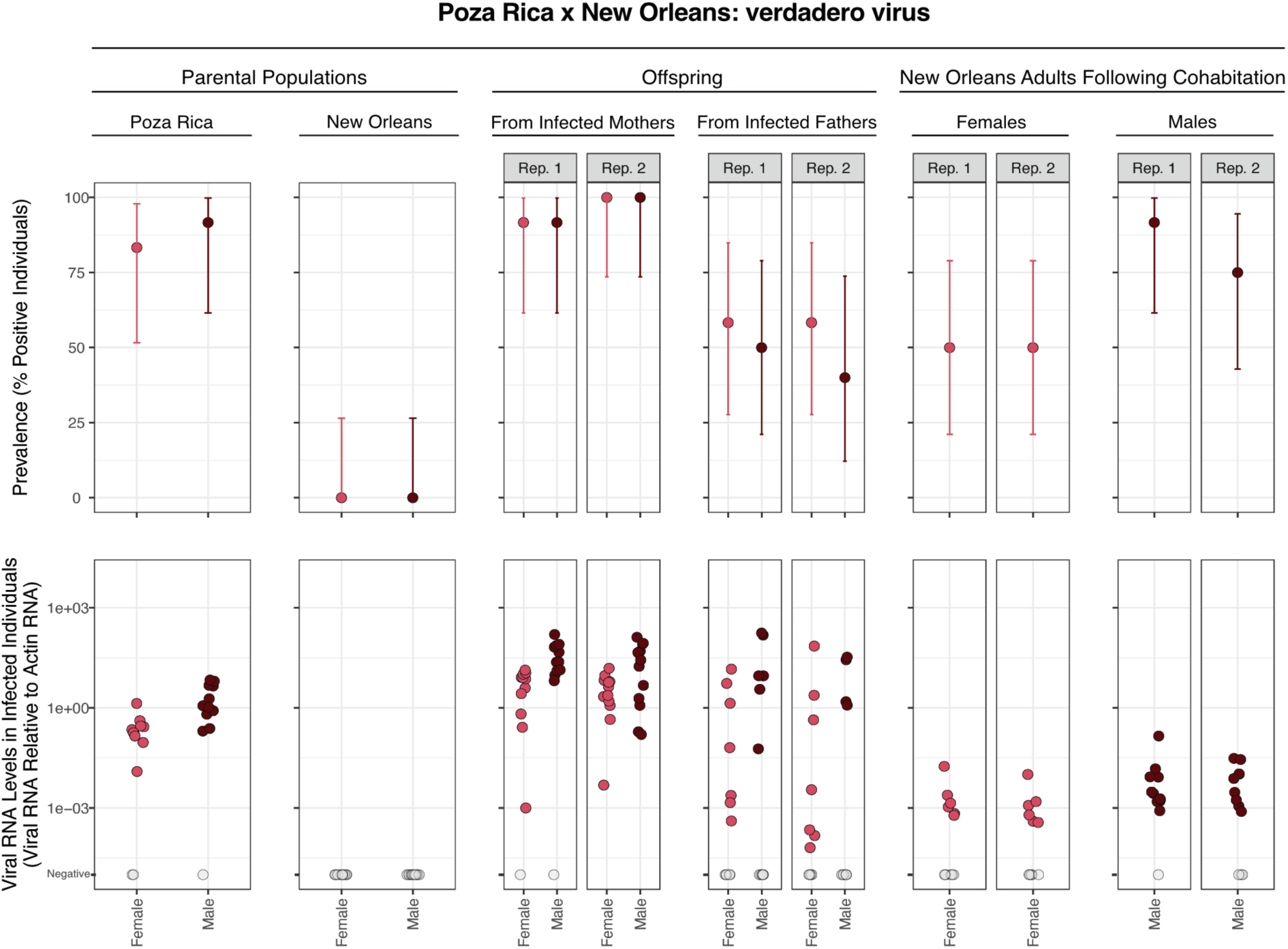
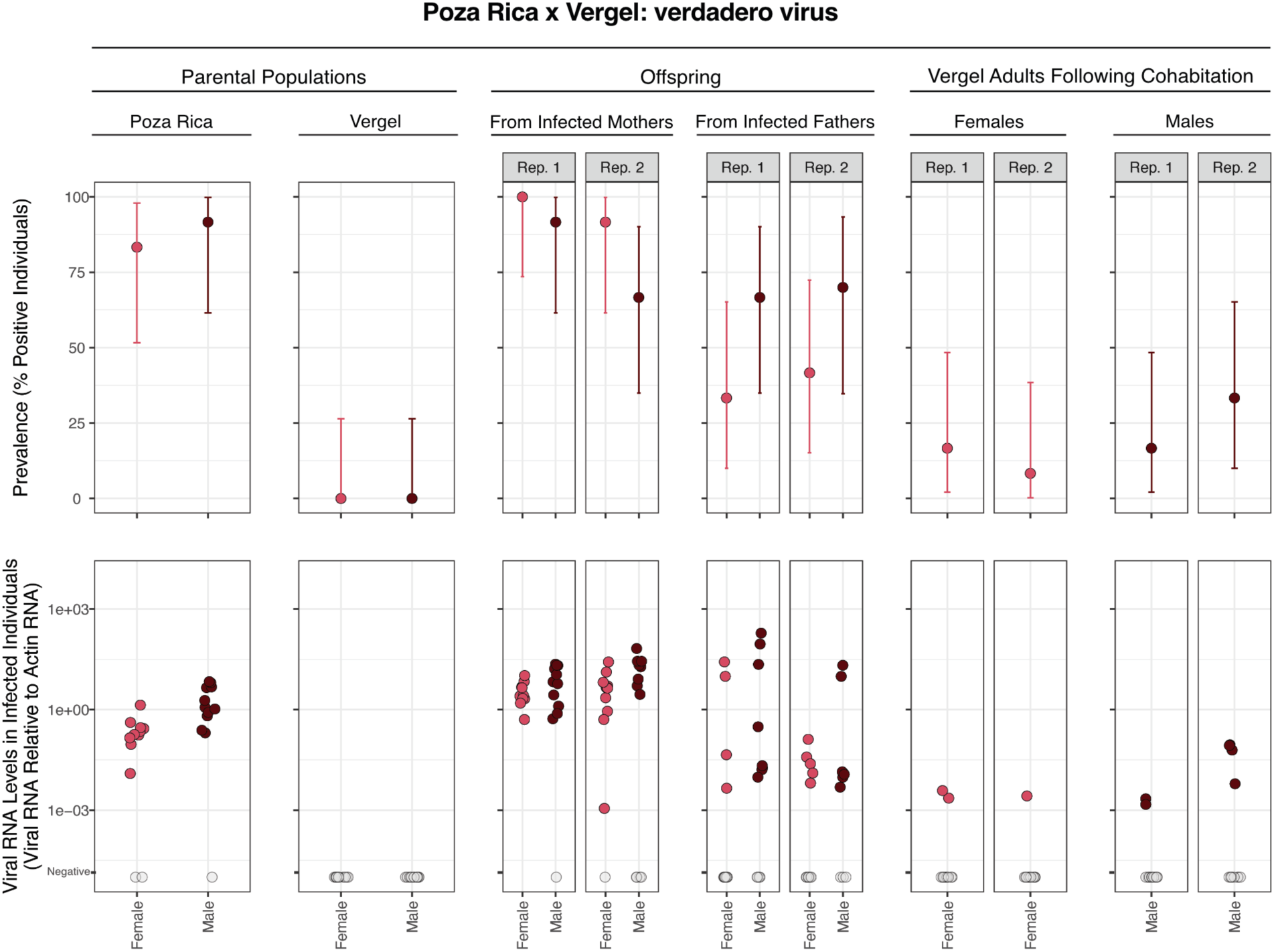
Prevalence and transmission of verdadero virus. Upper panels show prevalence of verdadero virus in different populations of parental and offspring mosquitoes. Error bars indicate binomial 95% confidence intervals. Lower panels show levels of verdadero virus RNA detected by RT-qPCR in individual mosquitoes normalized to levels of actin mRNA. Values for uninfected mosquitoes are plotted in grey on the lowest Y axis position. Rep refers to replicate crosses. (A) Crosses between mosquitoes from the Poza Rica colony and the New Orleans colony. (B) Crosses between mosquitoes from the Poza Rica colony and the Vergel colony. The same data for parental Poza Rica is shown in upper and lower panels.

Like anphevirus, verdadero virus exhibited biparental vertical transmission (**Figure 2**). 88/96 (92%) of the offspring of Poza Rica mothers tested positive for verdadero virus RNA. Paternal transmission was less efficient, with only 48/92 (52%) of the offspring of Poza Rica fathers. The mean level of anphevirus RNA in infected offspring was similar for maternal and paternal transmission: 17.0× and 18.8× the level of actin mRNA, respectively. Despite these similar averages, levels of verdadero virus RNA in individual offspring spanned a broad range, from 1.0×10^−5^ to 193× the levels of actin mRNA.

Some New Orleans and Vergel parents tested positive for verdadero virus RNA following cohabitation with opposite-sex Poza Rica parents (**Figure 2**). 32/48 (67%) of New Orleans parents and 9/48 (19%) of Vergel parents were positive following cohabitation (**Figure 2**). Verdadero virus RNA levels in these mosquitoes were low, on average 0.02× levels of actin mRNA.

We modeled verdadero virus transmission as we had for AeAV (**Table 3**). As with anphevirus, paternal vs maternal transmission was the only variable significantly predicting verdadero virus prevalance in offspring (**Table 3**), with offspring more likely to be infected via maternal transmission (p = 5.0×10^−8^). In contrast to AeAV, male adults were more likely to test positive for verdadero virus following exposure to infected Poza Rica females (p = 1.2×10^−2^). Adults from the New Orleans colony were more likely to test positive than those from the Vergel colony following cohabitation, indicating a possible role for mosquito genotype or behavior in susceptibility to verdadero virus infection (p = 6.2×10^−6^; **Figure 2**; **Table 3**).

**Table 3:** Logistic regression model of verdadero virus transmission.

| Predictor | Vertical transmission |  |  | Horizontal transmission |  |  |
| --- | --- | --- | --- | --- | --- | --- |
|  | OR | 95% CI | p-value | OR | 95% CI | p-value |
| (Intercept) | 12.6 | 5.46, 33.4 | <0.001 | 1.12 | 0.53, 2.38 | 0.8 |
| Offspring or exposed parent sex [male] | 1.01 | 0.49, 2.06 | >0.9 | 3.65 | 1.38, 10.6 | **0.012** |
| Non-infected parent colony [Vergel] | 0.77 | 0.37, 1.57 | 0.5 | 0.09 | 0.03, 0.25 | **<0.001** |
| Vertical transmission mode [paternal] | 0.10 | 0.04, 0.22 | **<0.001** |  |  |  |
| No. Obs. | 188 |  |  | 96 |  |  |
| AIC | 189.9 |  |  | 106.5 |  |  |

Abbreviations: CI = Confidence Interval, OR = Odds Ratio

### Guadeloupe mosquito virus

Guadeloupe mosquito virus (GMV) is a two-segmented positive sense single-stranded (ss) RNA virus similar to viruses in the *Solemoviridae* family (17, 18, 55–58). GMV has been detected in *Aedes* and other mosquito species and appears to have a global distribution. GMV was detected in both the Poza Rica and Tapachula colonies by metagenomic sequencing (**Table 1**).

All tested Poza Rica adults were positive for GMV RNA, and RNA levels were very high in individual mosquitoes: 3174.4× the level of actin mRNA on average (**Figure 3**). Tapachula adults were also mostly infected: 20/24 (83%) tested positive (**Supplemental Figure 4**). But GMV RNA levels were more variable in the Tapachula adults, with substantially lower levels in some positive mosquitoes (**Supplemental Figure 4**). Unexpectedly, some New Orleans and Vergel adults tested positive for GMV: 15/47 (32%) and 32/48 (67%), respectively (**Figure 3**). This was unexpected because no GMV-mapping reads were detected by metagenomics in these colonies. The lack of detection of GMV in these colonies by metagenomic likely reflects the low GMV RNA levels in infected mosquitoes (**Figure 3**). We confirmed the identity of the GMV PCR product amplified from New Orleans and Vergel adults by agarose gel electrophoresis and Sanger sequencing. The GMV sequences from the 4 colonies shared > 98.9% pairwise identity in the amplified region, but were not identical (**Supplemental Figure 5**). We also performed qPCR without reverse transcription to confirm that low-level detection did not result from transcription of a GMV-like endogenized viral sequence (50, 51). The presence of GMV in all parental populations meant that it was not possible to definitively determine whether infection in offspring derived from maternal or paternal transmission. However, the much more highly infected Poza Rica and Tapachula parents were the likely source of infection.

**Figure 3:**
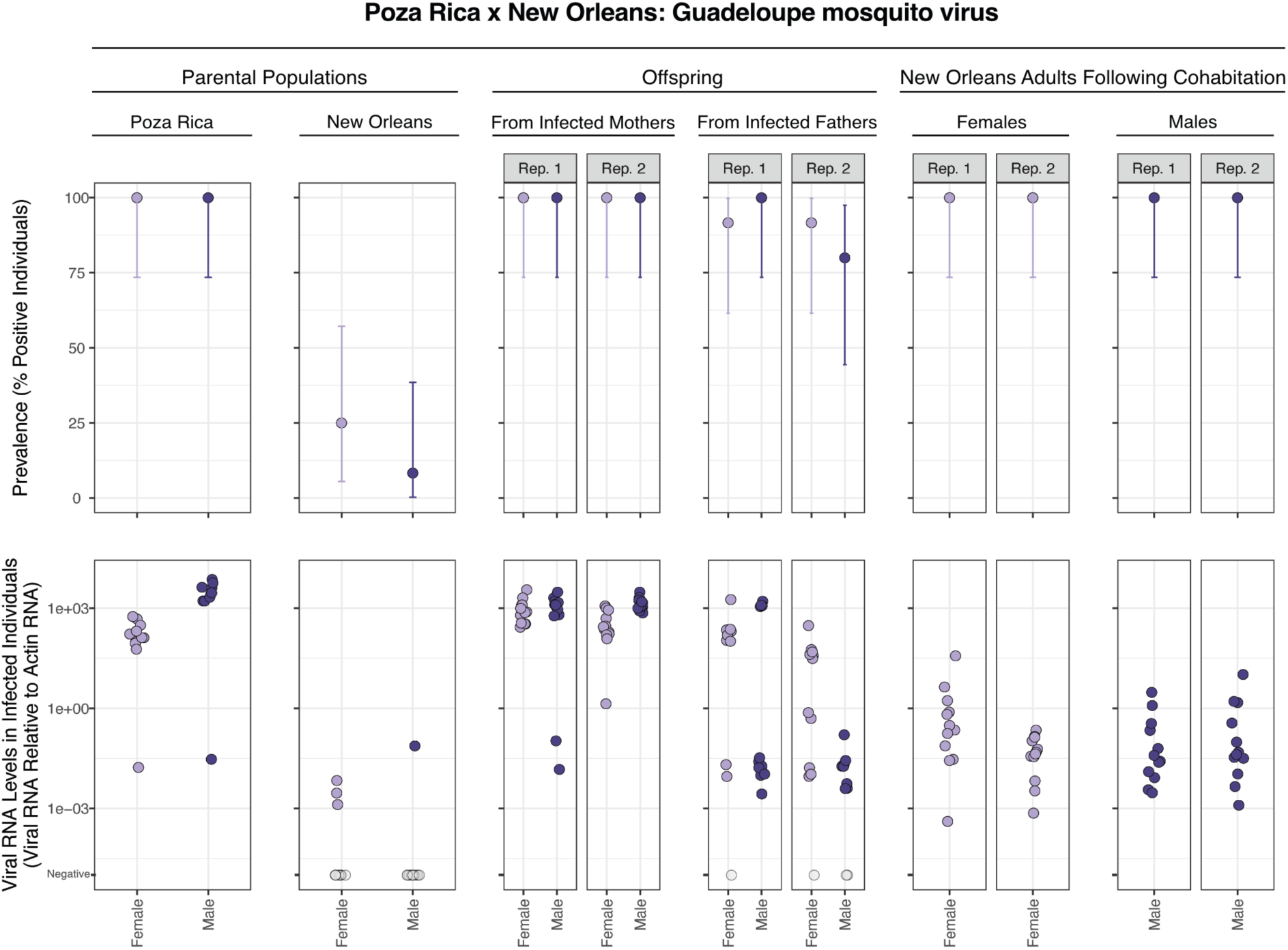
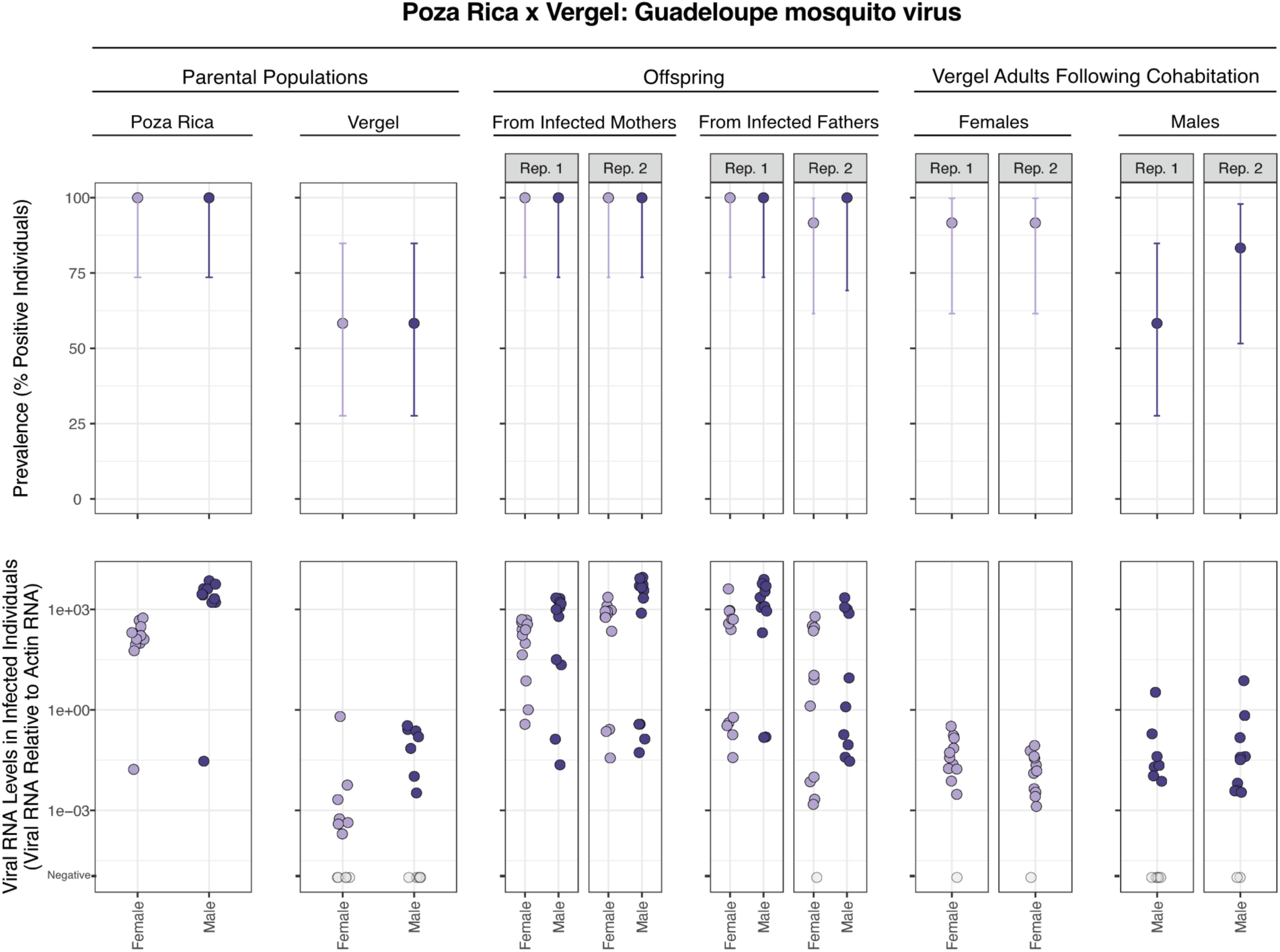
Prevalence and transmission of Guadeloupe mosquito virus from crosses involving Poza Rica parents. Upper panels show prevalence of GMV in different populations of parental and offspring mosquitoes. Error bars indicate binomial 95% confidence intervals. Lower panels show levels of GMV RNA detected by RT-qPCR in individual mosquitoes normalized to levels of actin mRNA. Values for uninfected mosquitoes are plotted in grey on the lowest Y axis position. Rep refers to replicate crosses. (A) Crosses between mosquitoes from the Poza Rica colony and the New Orleans colony. (B) Crosses between mosquitoes from the Poza Rica colony and the Vergel colony. The same data for parental Poza Rica is shown in upper and lower panels. Infected mothers and infected fathers refers to the sex of the much more highly infected parents from the Poza Rica population.

Transmission of GMV from infected Poza Rica parents was efficient. 96/96 (100%) and 87/92 (95%) of offspring of Poza Rica mothers and fathers tested positive for GMV RNA. In both cases infected offspring exhibited a wide range of GMV RNA levels (**Figure 3**). GMV was at lower prevalence and RNA levels in parental Tapachula mosquitoes, and, accordingly, fewer offspring of Tapachula parents were infected. 88/96 (92%) and 42/92 (46%) offspring of Tapachula mothers and fathers were positive for GMV, respectively (**Supplemental Figure 4**).

We modeled GMV transmission as we had for other viruses (**Table 4**). GMV models included an additional term corresponding to the highly-infected parental colony: Poza Rica vs. Tapachula. This term captured possible differences in parental host genetics and differences in starting prevalence and viral loads between these colonies (**Figure 3**, **Supplemental Figure 4**). In the vertical transmission model, transmission from highly infected mothers was more efficient (p = 2.7×10^−10^), and transmission from Poza Rica crosses was more efficient than Tapachula crosses, likely reflecting higher GMV starting prevalence and RNA levels in the Poza Rica population (p = 2.5×10^−10^).

**Table 4:** Logistic regression model of GMV transmission. Main infected parent colony refers to the highly-infected parental colony: Poza Rica vs. Tapachula. Less infected parent colony reflects the parental colony with low-level infection: New Orleans vs. Vergel. The horizontal transmission model included replicate as a random effect.

| Predictor | Vertical transmission |  |  | Horizontal transmission |  |  |
| --- | --- | --- | --- | --- | --- | --- |
|  | OR | 95% CI | p-value | OR | 95% CI | p-value |
| (Intercept) | 194 | 62.4, 741 | <0.001 | 71.8 | 10.4, 496 | <0.001 |
| Offspring or exposed parent sex [male] | 1.17 | 0.59, 2.30 | 0.7 | 0.13 | 0.03, 0.58 | **0.007** |
| Main infected parent colony [Tapachula] | 0.04 | 0.01, 0.10 | **<0.001** | 0.02 | 0.00, 0.10 | **<0.001** |
| Less infected parent colony [Vergel] | 1.91 | 0.97, 3.83 | 0.063 | 0.60 | 0.13, 2.68 | 0.5 |
| Vertical transmission mode [paternal] | 0.07 | 0.03, 0.15 | **<0.001** |  |  |  |
| No. Obs. | 376 |  |  | 191 |  |  |
| AIC | 227.8 |  |  | 160.5 |  |  |

Abbreviations: CI = Confidence Interval, OR = Odds Ratio

The impact of cohabitation and mating on GMV prevalence in New Orleans and Vergel adults was variable (**Figure 3**, **Supplemental Figure 4**). The GMV horizontal transmission model that included replicate as a random effect was better fitting than the fixed effect only model, reflecting increased variance in prevalence between replicate crosses (**Figure 3**, **Supplemental Figure 4**). New Orleans and Vergel females were more likely to test positive for GMV following cohabitation, again reflecting sexual transmission as a candidate route of exposure (**Table 4**; p = 6.8×10^−3^). Cohabiting mosquitoes were also more likely to test positive following exposure to Poza Rica adults (**Table 4**; p = 1.3×10^−6^).

**Supplemental Figure 4:**
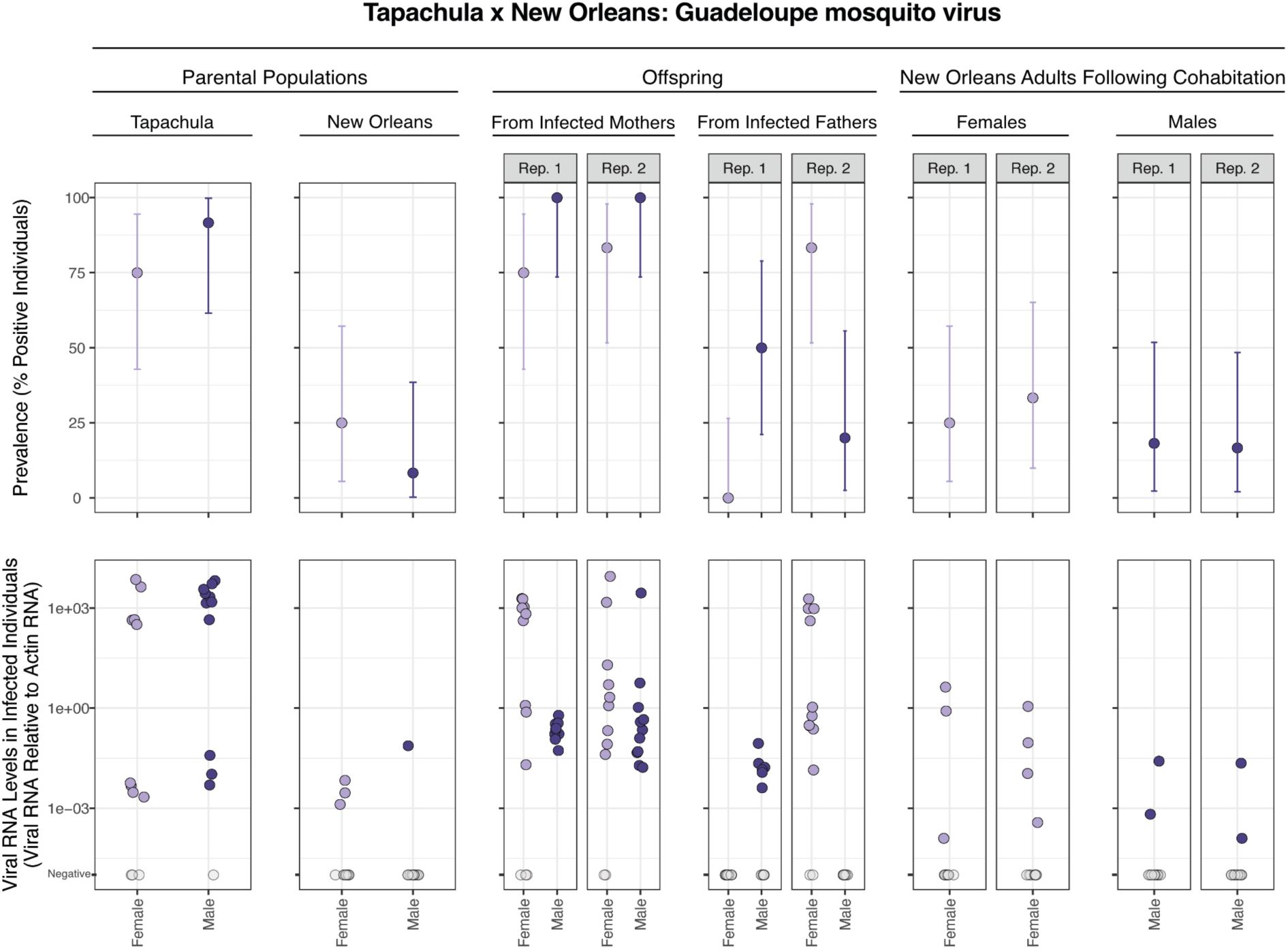
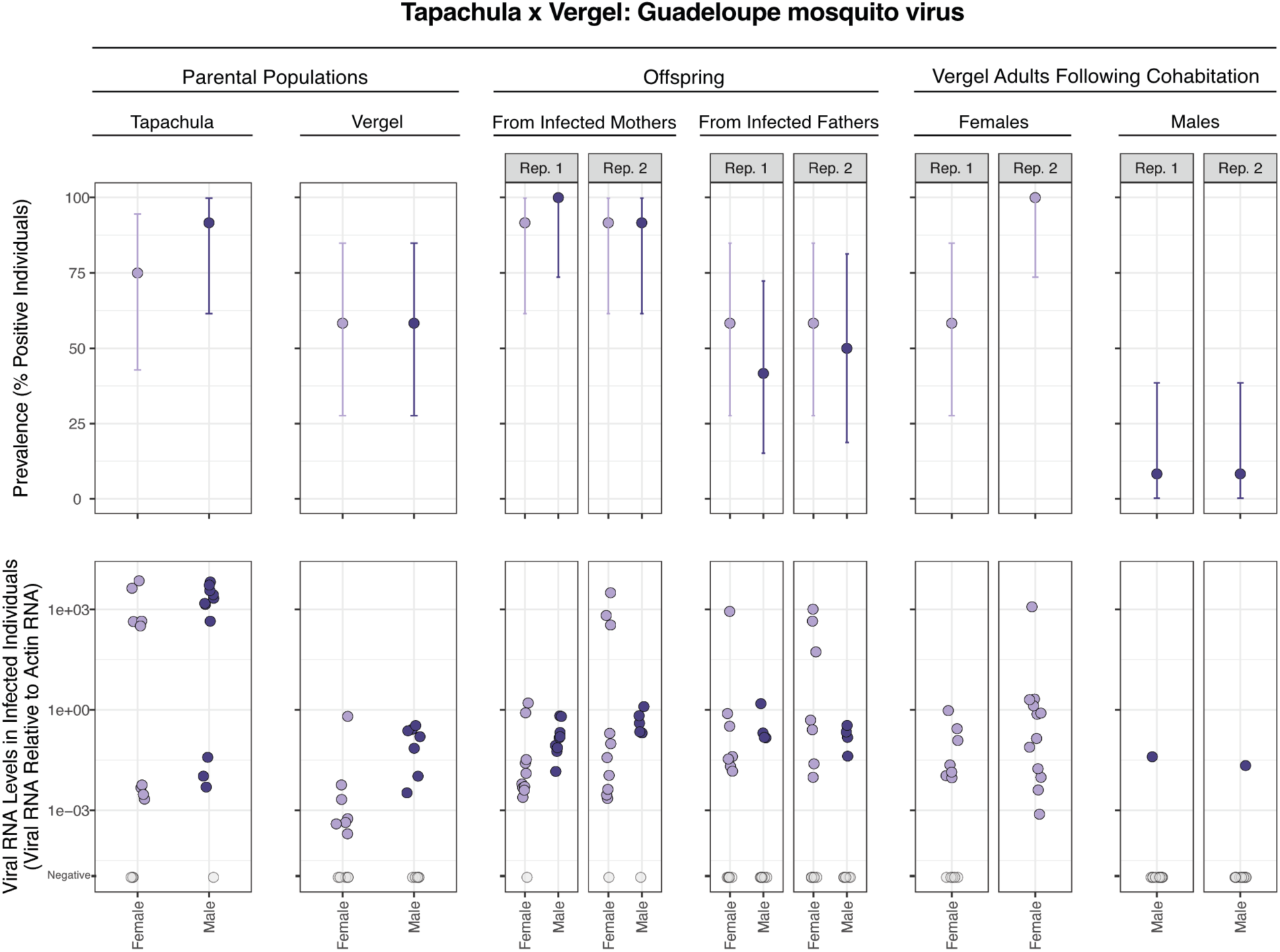
Prevalence and transmission of Guadeloupe mosquito virus from crosses involving Tapachula colony parents. Upper panels show prevalence of GMV in different populations of parental and offspring mosquitoes. Error bars indicate binomial 95% confidence intervals. Lower panels show levels of GMV RNA detected by RT-qPCR in individual mosquitoes normalized to levels of actin mRNA. Values for uninfected mosquitoes are plotted in grey on the lowest Y axis position. Rep refers to replicate crosses. (A) Crosses between mosquitoes from the Tapachula colony and the New Orleans colony. (B) Crosses between mosquitoes from the Tapachula colony and the Vergel colony. Data for parental New Orleans and Vergel mosquitoes as in Figure 3. The same data for parental Tapachula is shown in upper and lower panels. Infected mothers and infected fathers refers to the sex of the much more highly infected parents from the Poza Rica population.

## Discussion

In this study, we characterized the viromes of four long-established colonies of *Aedes aegypti* and identified three prevalent insect-specific viruses: *Aedes* anphevirus, verdadero virus, and Guadeloupe mosquito virus. Their persistent high prevalence in these populations is consistent with minimal fitness costs and an efficient ability to transmit between generations.

We measured two types of possible transmission for these viruses: vertical transmission across generations, and horizontal transmission between cohabiting and mating adults. In both cases, we did not assess exact mechanisms of transmission, and multiple mechanisms may contribute. Transmission to offspring from infected mothers may occur via infected eggs (9, 59).

Alternatively, the exterior of eggs may harbor infectious virus particles that could infect larvae following hatching. Horizontal transmission between aquatic larvae or pupae is also possible. Regardless of the exact transmission route(s), maternal transmission of all three viruses was efficient, with all or nearly all adult offspring infected. Transmission from infected fathers was uniformly less efficient and could involve infectious virus particles in or on sperm or in seminal fluid (60–62).

There were several limitations to this study. We did not perform single-pair crosses, so did not measure the infection status of individual parents (63). Some of the uninfected offspring may simply have had uninfected parents. However, parental prevalences were at or near 100% (**Figure 1**, **Figure 2**, **Figure 3**), and the large numbers of parents in crosses meant that offspring provided population-level estimates of transmission efficiency. We also did not determine the extent to which low level RT-qPCR signals reflected genuine low level infections rather than contaminating RNA acquired during cohabitation with infected mosquitoes. We tracked infection dynamics across a single generation of offspring. It is possible that, as with *Drosophila melanogaster* sigmavirus, offspring infected paternally are infected but less capable of further vertical transmission (64, 65).

Although paternal transmission was less efficient than maternal transmission, it still might contribute to the maintenance of these viruses in natural populations (66). Biparentally-transmitted symbionts have the ability to increase in prevalence even if infection reduces host fitness. In contrast, vertically transmitted symbionts with maternal-only transmission are expected to decline in prevalence if there is any fitness cost from infection (66). This is why organisms with maternal-only transmission, like *Wolbachia* endosymbiotic bacteria, have frequently evolved mechanisms to manipulate host reproduction to maintain their prevalence in subsequent generations (67). Biparentally-transmitted viruses such as those studied here can persist through vertical transmission alone, provided that the combined maternal and paternal transmission efficiency exceeds 100% and that infection is not too costly (65, 66). Biparental vertical transmission therefore represents a valuable capability and a means to drive symbionts to high prevalence in host populations, as was evident here. However, it is unclear why GMV was at lower prevalence and RNA levels in the New Orleans and Vergel populations. These lower-level infections may reflect differences in host or viral genetics, although the partial GMV sequences from the four colonies were nearly identical (Supplemental Figure 5).

Horizontal transmission can also contribute to maintenance of inherited symbionts in host populations (66). Increases in prevalence in New Orleans and Vergel mosquitoes following cohabitation were consistent with possible horizontal transmission, although RNA levels in individual positive adults were much lower than levels in infected offspring. Higher AeAV and GMV prevalences in exposed females following mating are consistent with possible sexual transmission. Whether possible horizontal transmission of these viruses contributes meaningfully to overall prevalence in a background of robust vertical transmission remains to be determined (66).

A major motivation for studying ISVs has been their potential utility in mosquito control and population modification. Some ISVs may naturally reduce vector competence, analogous to the effects of certain strains of *Wolbachia* (68). Alternatively, engineered derivatives of these viruses could be used to deliver transgenes to mosquito populations (21, 22). One reason *Wolbachia*-based strategies have been successful is that the endosymbiont can spread through mosquito populations following limited initial release (69). CRISPR-based gene drives represent another example of a self-propagating approach to mosquito control (70). Viruses with an inherent capacity to spread via biparental vertical transmission, such as *Aedes* anphevirus, Guadeloupe mosquito virus, and verdadero virus, therefore represent promising candidates for vector control applications. The development of reverse genetic systems for these and other ISVs will be critical for determining their amenability to genetic modification and for evaluating their potential as scalable tools for mosquito control.

## Acknowledgements

The authors wish to thank Kai Chase, Susi Bennett, Irma Sanchez-Vargas, Megan Miller, Ashley Janich, Greg Pugh, and Bill Black IV, posthumously.

## Supplemental Material

**Supplemental Table 1:**
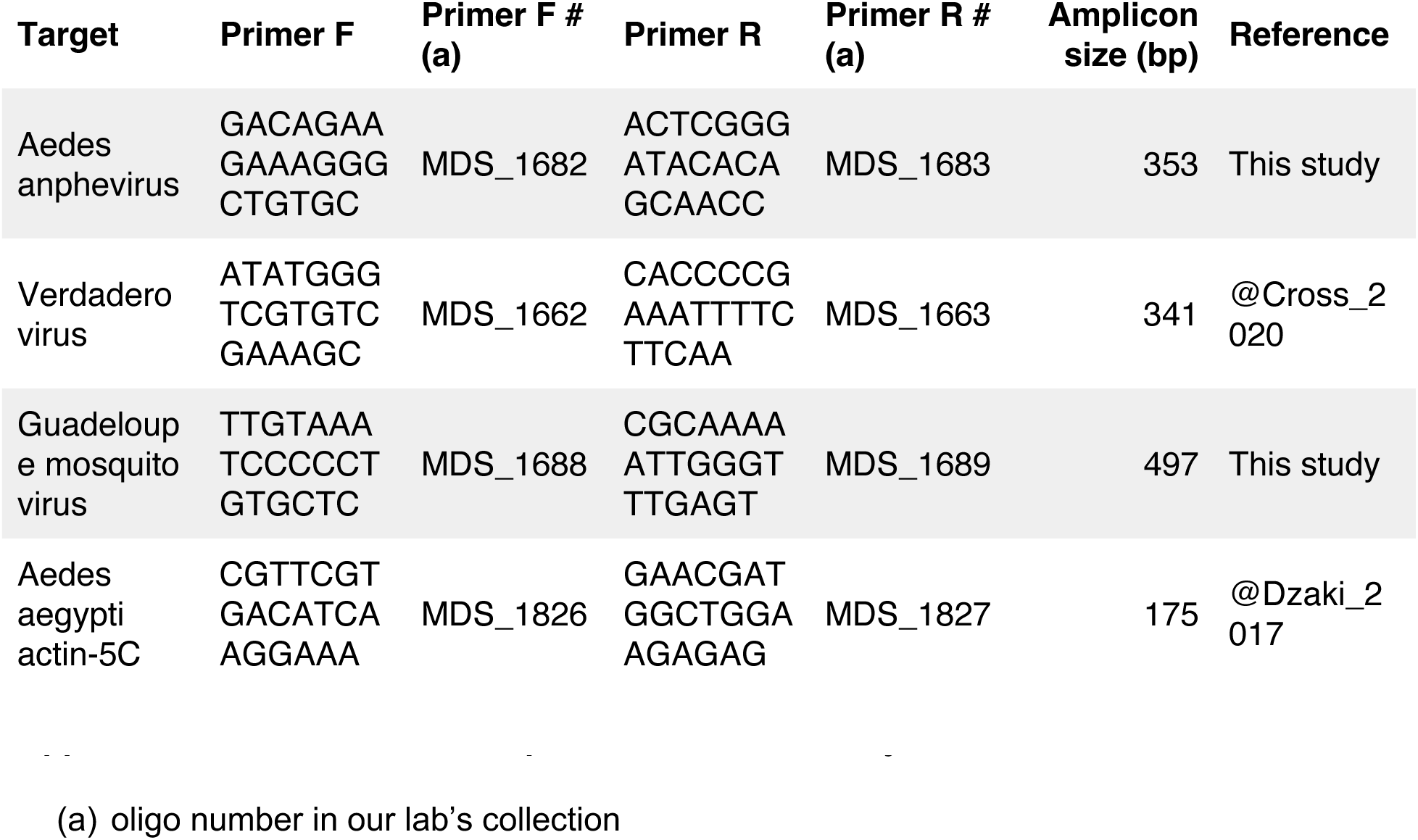
Primer sequences used this study.

**Supplemental Figure 5:**
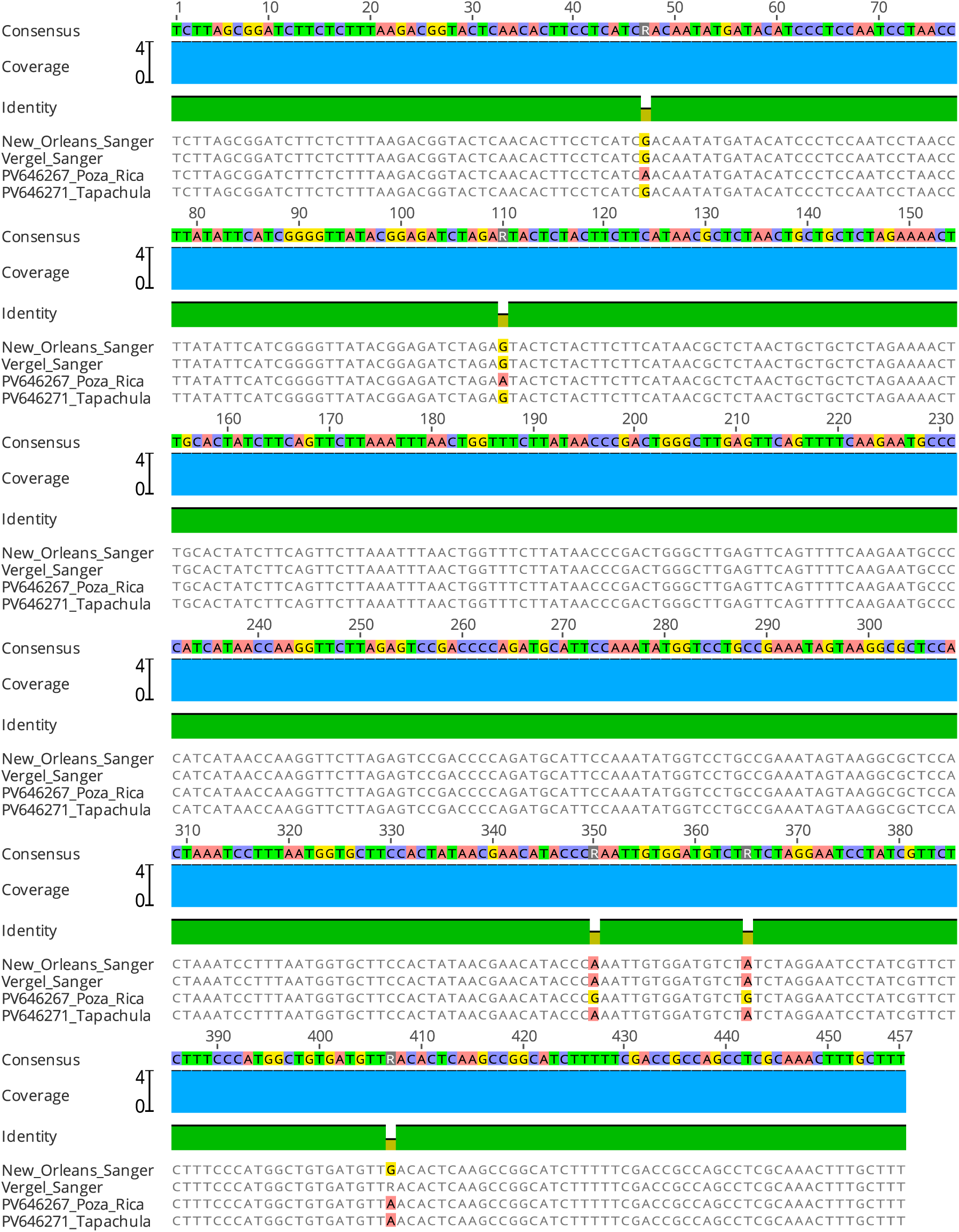
A multiple sequence alignment of Guadeloupe mosquito virus RNA 1 sequences from the four Aedes aegypti colonies. This alignment shows the region amplified by the primers used to detect and quantify GMV, excluding primer-binding sites Supplemental Table 1. The sequences from the New Orleans and Vergel colonies were determined using Sanger sequencing. The sequences from the Poza Rica and Vergel colonies were determined using metagenomic RNA sequencing. An R base in the Vergel sequence corresponds to a position where the Sanger chromatogram showed equal-height A and G peaks.

